# Early signs of plant community responses to climate warming along mountain roads in Switzerland

**DOI:** 10.1101/2024.07.12.603213

**Authors:** Evelin Iseli, Nathan Diaz Zeugin, Camille Brioschi, Jake Alexander, Jonathan Lenoir

## Abstract

1. Global warming is pushing species to shift their ranges towards higher latitudes and elevations, causing a reassembly of plant communities potentially accompanied by community thermophilization (i.e., an increasing number or cover of thermophilic species, sometimes at the expense of mesic or cold-adapted species). Given the large variation typically observed in the magnitude and direction of range shifts, quantifying community thermophilization might provide a sensitive method to detect range shifts within short time periods and across limited spatial extents. Assessing changes in plant community composition as a whole might integrate early signs of range shifts across co-occurring species while accounting for changes in both occurrence and abundance.
2. Here, we combine an assessment of (i) species-level range shifts, (ii) changes in species richness and (iii) changes in community-inferred temperatures along three mountain roads in Switzerland to ask whether plant communities have responded to warming climate over a 10 year period, and whether community thermophilization is an appropriate metric for early detection of these changes.
3. We found a community thermophilization signal of +0.13°C over the 10-year study period based on presence-absence data only. Despite significant upward shifts of species’ upper range limits in the lower part of the studied elevational gradient and a decrease in species richness at high elevations, significant thermophilization was not detectable if community- inferred temperatures were weighted by species’ covers. Low cover values of species that were gained or lost over the study period and their similar species-specific temperatures to resident species explained the discrepancy between the thermophilization detected in either cover-weighted or unweighted models.
4. *Synthesis.* Our work shows that plant species are shifting to higher elevations along roadsides in the western Swiss Alps and that this translates into a detectable warming signal of plant communities within 10 years. However, the species-level range shifts and the community-level warming effect are mostly based on low cover values of gained/lost species, preventing the detection of community thermophilization signals when incorporating cover changes. We therefore recommend using unweighted approaches for early detection of community-level responses to changing climate, ideally set into context by also assessing species-level range shifts.

## INTRODUCTION

Plant species around the world are shifting their geographical distributions along elevational and latitudinal gradients in response to changing climates (Lawlor et al., 2024; Lenoir & Svenning, 2015; Spence & Tingley, 2020). The dynamic nature of range expansions and retractions is leading to a reassembly of communities, potentially adding to the direct consequences of climate change such as biodiversity loss and gain, or altered ecosystem functions (Rumpf et al., 2019; Steinbauer et al., 2018). Due to elevation dependent warming, mountain ecosystems are especially exposed to global warming and thus affected by climate change (Pepin et al., 2015), which could be further exacerbated by rapid elevational range shifts as species can benefit from steep environmental gradients and correspondingly short dispersal distances (Alexander et al., 2018). Detecting range shifts and understanding their potential impacts is fundamental for making informed predictions regarding the influence of climate warming on biodiversity, ecosystem functioning and the associated provisioning of ecosystem services (Pecl et al., 2017; Sintayehu, 2018).

While average trends of upslope range shifts are often detected and interpreted as signals of climate change (Feeley et al., 2013; Rumpf et al., 2018; Rumpf et al., 2019; Steinbauer et al., 2018), there is substantial variation in the magnitude of range shifts between species, ecosystems and regions (Lenoir et al., 2020; Lenoir et al., 2010; O’Sullivan et al., 2021). Various studies have found that many species and taxonomic groups do not show any range shifts at all and that a large proportion of range shifts are reported to be in an unexpected direction (e.g., downwards shifts in the elevational context). In recent reviews, common range-shift expectations were only met in 47% to 59% of range shift observations (Lawlor et al., 2024; Rubenstein et al., 2023). This large variation in both the magnitude and direction of range shifts between species makes it difficult to detect significant signals of range shifts within short time periods and across limited spatial extents.

Range shift studies often also focus on a single range property only, such as the centroid or the upper elevational range limit (i.e., the leading edge), while changes of the lower elevational range limit (i.e., the trailing edge) as well as abundance changes within a species’ current range are often neglected (Cannone & Pignatti, 2014; Lawlor et al., 2024; Rubenstein et al., 2023; Rumpf et al., 2018; Rumpf et al., 2019). Only a joint evaluation of range expansion, retraction and abundance changes (i.e., a multidimensional assessment as suggested in Lenoir & Svenning, 2015) allows a comprehensive evaluation of the complex dynamics of climate change effects on species’ range dynamics, and so whether a particular species benefits or suffers from a changing climate. The importance of combining range shifts with abundance changes has been highlighted in a recent study on marine fish, which showed that contrary to the general assumption, rapid range shifts are not necessarily connected with increasing population abundances at the historical position of the leading edge and that fast-moving species might be as vulnerable to climate change as slow-moving species (Chaikin et al., 2024). Indeed, abundance increases at the leading edge within the current range (i.e., in-filling) can be an important response to a changing climate and even outweigh the effect of range-limit shifts in community changes (Cannone & Pignatti, 2014; Rumpf et al., 2018). However, abundance shifts are frequently ignored because the detection of abundance changes requires an increased sampling effort (i.e., occurrence data is not sufficient) or long time series.

Community thermophilization – a reshuffling of the species assemblage towards a composition more typical of a warmer region – is driven by the gain of newly arriving warm-adapted species, the loss of cold-adapted species and changing abundances of species already present in the community (Gottfried et al., 2012). Hence, thermophilization integrates species abundance changes at a community level as well as local colonization and extinction events associated with leading- and rear-edge shifts. Recent reports of community thermophilization have often been related to leading-edge upward shifts, colonization events and increasing species richness (Bertrand et al., 2011; Cuesta et al., 2023; Gottfried et al., 2012; Hamid et al., 2020; Kiebacher et al., 2023; Martin et al., 2019; Verma et al., 2023), while the contribution of rear-edge contractions, local extinctions and abundance changes are less often considered explicitly (but see Duque et al., 2015; Fadrique et al., 2018; Feeley et al., 2013; Hamid et al., 2020; Lamprecht et al., 2018; Martin et al., 2019). Studies which focus on elevational gradients frequently find varying thermophilization patterns involving different potential mechanisms at different elevations (Bertrand et al., 2011; Fadrique et al., 2018; Hamid et al., 2020; Kiebacher et al., 2023; Savage & Vellend, 2015; Verma et al., 2023). Both increasing (Bertrand et al., 2011; Kiebacher et al., 2023; Savage & Vellend, 2015; Verma et al., 2023) and decreasing (Fadrique et al., 2018; Hamid et al., 2020) thermophilization rates at higher elevation have been observed. Lower thermophilization rates at low elevation could be explained by a higher proportion of warm-adapted and cosmopolitan species and therefore greater tolerance of increasing temperatures, longer dispersal distances as well as dispersal barriers caused by habitat fragmentation (Bertrand et al., 2011; Kiebacher et al., 2023) and reduced variation in temperatures due to a forest habitat (Verma et al., 2023). Contradicting the general trend of increased warming at higher elevations, Fadrique et al. (2018) connect the lower thermophilization rate they reported at high elevations with reduced warming rates and the distinctively different ecotones along the Andean slopes.

Given the lack of studies combining all of the suggested explanations for reported community thermophilization patterns and their variation along elevational gradients (range shifts of the leading- and rear-edge, local colonizations and extinctions and abundance changes), a combined analysis of the proposed mechanisms is needed. Such an analysis could not only shed light on the relationship between range shifts and abundance changes and how they combine to create changes at the community level but it could also help identify sensitive ways to detect community responses to climate change over short time periods. In this study, we analysed plant community thermophilization rates along mountain roads in the southwestern Swiss Alps. Compared to other long-term vegetation monitoring initiatives, our dataset – based on the Mountain Invasion Research Network (MIREN; www.mountaininvasions.org; Haider et al., 2022) protocol – is specifically designed to detect and study range shifts and community composition changes along elevational gradients. The high density of survey plots along few chosen mountain roads allows a focused assessment of local species distributions within comparable environmental conditions and the efficient survey design allows a high temporal resolution compared to more extensive national surveys (e.g. BDM in Switzerland; www.biodiversitymonitoring.ch) while keeping costs relatively low. We also assessed which processes (gain of new warm-adapted species, loss of cold-adapted species or changes in abundances) are responsible for the observed changes in community-inferred temperatures. By comparing thermophilization rates at the community level to species-level range shifts, we discuss the relative merits of these two complementary approaches to improve our understanding of species redistributions. Specifically, we ask: (1) Have community-inferred temperatures increased over the study period of 10 years and do rates of change vary along the studied elevational gradient? (2) To what extent are community-inferred temperature changes driven by the gain vs. loss of species vs. abundance changes within the existing community? (3) Are community-level or species-level assessments of climate change impacts more sensitive over short time periods? We hypothesise that a multidimensional analysis of species distribution and abundance changes yield a clearer signal of climate change impacts on plant assemblages when high variation between species and short study periods constrain the detection of species-level range shifts (Katabuchi et al., 2017).

## METHODS

### Study sites and vegetation data

Our analysis of community thermophilization is based on two independent data sets, the Mountain Invasion Research Network data (MIREN data, the data set of interest) and the Swiss Biodiversity Monitoring data (BDM data, the training data set). The MIREN data (Haider et al., 2022) consist of time series of vegetation surveys along mountain roads around the world, with the first regions surveyed for the first time in 2007-2008 and repeated in 5-year intervals. In each of the participating regions, three roads were selected and evenly stratified into 20 transects between the top (highest point of the road) and bottom (point below which no major elevational change occurs). All roads are open to traffic for at least part of the year. Each transect is composed of three rectangular 2 m × 50 m plots placed in a T-shape along the focal road, with one plot parallel to the road and two aligned and contiguous plots perpendicular to it to differentiate between disturbed habitats on the road verge and less disturbed habitat away from the road verge. The two perpendicular plots were only surveyed in the absence of major barriers or cliffs. Vegetation surveys include recording the identity of all vascular plants and estimating their cover according to a standard protocol (Haider et al., 2022; information on cover classes in Table S1). For this study, only the Swiss MIREN data were considered (Figure 1), which consist of three roads, 63 T-shape transects and 137 plots in the cantons of Vaud and Valais in the southwestern Swiss Alps. The mean annual temperature in 2022, the year of the last survey, ranged from 4.04°C at 1800 m a.s.l. at the top of the studied elevational gradient near Brig, Switzerland to 12.6°C at 415 m a.s.l. at the bottom of the gradient near Bex, Switzerland.

**Figure 1.**
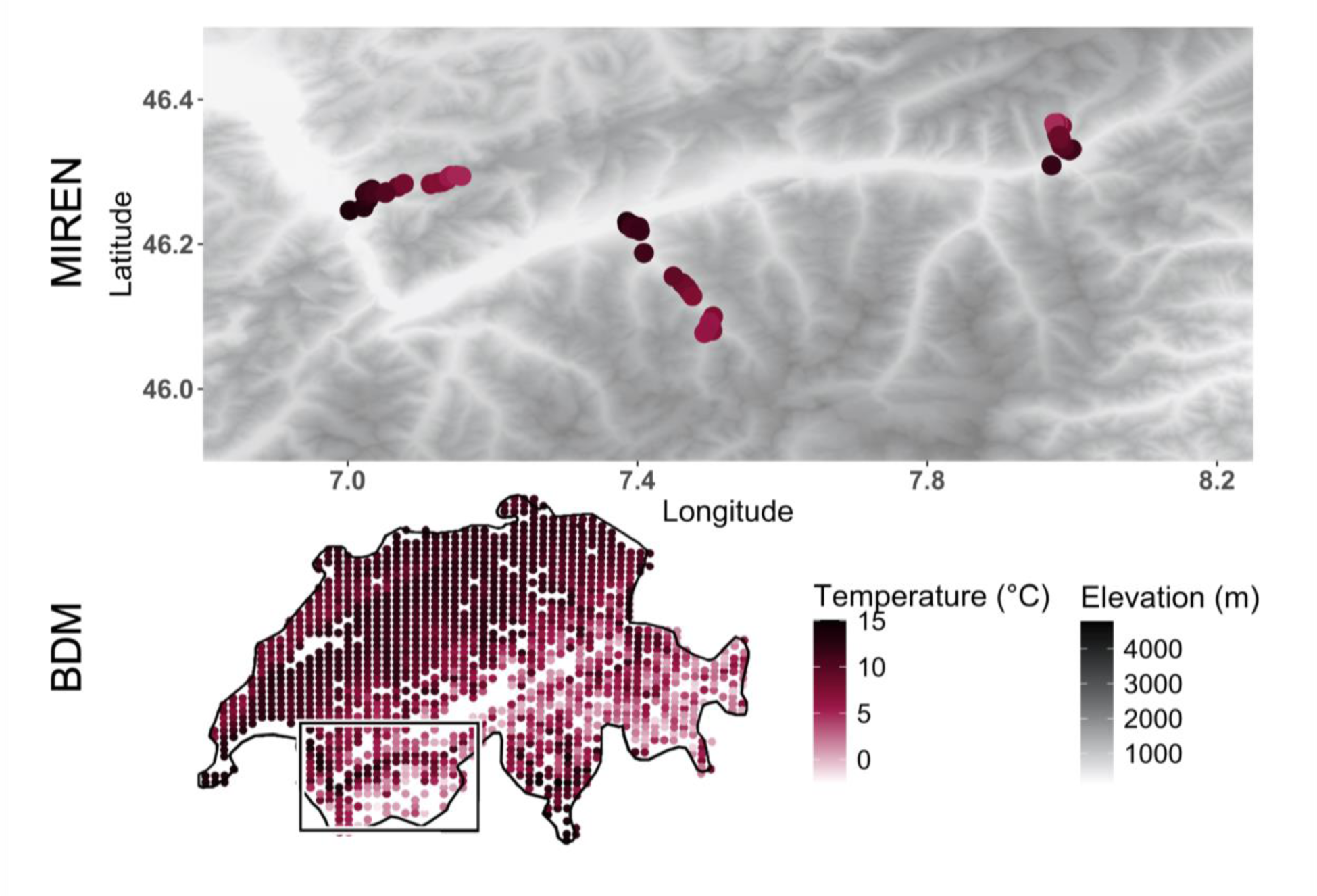
Distribution of survey plots of training data set (BDM, lower panel) and data set of interest (MIREN, upper panel) in Switzerland. Red points in the upper panel indicate the location of the 62 repeatedly surveyed Swiss MIREN transects along three mountain roads, red points in the lower panel mark the location of all 1161 vegetation survey plots of the Swiss Biodiversity Monitoring data used to train the fxWAPLS model. The red colour gradient depicts the mean annual temperature (rolling average over five years) in 2022 of each plot. The gray colour gradient shows the elevation of the area surrounding the MIREN transects in the upper panel. The location of the upper panel within Switzerland is indicated with a black frame.

The Swiss Biodiversity Monitoring data (BDM Coordination Office, 2014) provide a time series of vegetation surveys of 1448 circular 10 m^2^ permanent plots placed in a regular grid over Switzerland (Figure 1). The mean annual temperatures in 2022 range from -3.34°C near Zermatt, Switzerland to +14.37°C in Lavertezzo, near Locarno, Switzerland. The initial surveys were conducted between 2001 and 2005 and all plots are regularly re-surveyed every 5 years, but not all in the same year. A survey year consists of two visits to each of the plots (except for alpine habitats, which are only visited once), one in the beginning of the growing season and one at peak vegetation. During each visit, all vascular plants are identified and their cover estimated using a simplified Braun-Blanquet scale (Braun-Blanquet, 1932; for details on cover classes see Table S1) according to a standardized protocol (Auftragnehmer Biodiversitäts- Monitoring Schweiz, 2020).

To harmonize the taxonomy and format of the two data sets, the standardized MIREN species names (recorded according to WorldFlora online, http://www.worldfloraonline.org) were transformed into official Swiss taxonomy (https://www.infoflora.ch/en/flora/taxonomy.html) and aligned to BDM nomenclature based on both taxonomies. Non-matching MIREN species were manually searched, checked for synonyms within the BDM nomenclature and excluded from the MIREN data if not present in the BDM data at all. Following this exclusion, the MIREN data consisted of a total of 759 and the BDM data of a total of 1372 vascular plant species. To align the differing cover classes of the two datasets, the midpoint of the respective cover classes was adapted as the standard cover estimate (Table S1). As our analysis depends on a quantitative measure of the frequency of a species within a community, surveys with missing cover estimates were deleted. Cover was not recorded in MIREN surveys in 2007 at all and in some plots incompletely in the years after, reducing the number of surveys included in the analysis from 488 (137 plots × 4 sampling timepoints at the maximum: 2007, 2012, 2017, 2022) to 366 (122 plots × 3 sampling time points: 2012, 2017 and 2022). Using the oldest survey of each plot with cover estimates — similarly to MIREN, cover was not recorded in the earliest BDM campaigns — the BDM data set was reduced to one survey per plot to avoid pseudo-replication when training the model. Additionally, the two annual visits per survey year for the BDM data were combined into one by only retaining the larger cover estimate of a given plant species for that specific year. After cleaning, the resulting BDM dataset contained 1387 surveys (1 time point of 1387 plots) recorded between 2006 and 2015.

### Climate data

To obtain recent climate data (up to 2022) at a high spatiotemporal resolution, time series of mean annual temperature at a yearly timestep and at 1 km resolution across Switzerland were calculated following the change factor methodology (Anandhi et al., 2011). First, both CHELSAcruts monthly minimum and maximum temperatures — 30 arcsec resolution, 1902- 2016 (Karger et al., 2017; Karger & Zimmermann, 2018) — and TerraClimate monthly minimum and maximum temperatures — 150 arcsec resolution, 1958-2022 (Abatzoglou et al., 2018) — were each aggregated to mean annual temperatures. Then, a transformation factor between each smaller CHELSAcruts pixel within the bigger TerraClimate pixel was calculated for the time period from 1995 to 2015. We assumed that the anomalies computed for the overlapping period are consistent for the period after 2015 and used the calculated transformation factor to downscale TerraClimate data to CHELSAcruts resolution (30 arcsec resolution, which is about 1 km at the equator) from 2001 to 2022. Temperature values were integrated over five-year periods to smooth short-term fluctuations. The climate data were then used to extract the mean annual temperature of the survey year for all vegetation surveys used in the complete analysis (1 time point for the original 1448 BDM plots and 3 timepoints for the original 63 MIREN transects). As MIREN plots within a transect are located too close to each other for the resolution of the climate data, mean annual temperature for MIREN surveys was extracted at the transect level, using the mean of all plot coordinates per transect.

### Statistical analysis

To infer community-level plot temperature for MIREN plots along the studied elevational gradient, we used the statistical approach of fx-corrected tolerance-weighted weighted averaging partial least squares regression (fxTWAPLS). The method is often used to reconstruct paleoclimates from modern pollen assemblages but is useful for calculating any community-inferred temperatures (CIT) based on an independent training data set. The fxTWAPLS regression approach is an improved version of weighted averaging partial least square regression (WAPLS; ter Braak & Juggins, 1993), designed to reduce the compression towards the centre of the climatic gradient of the training data set. It incorporates a tolerance- weighting parameter to make use of the fact that species with a narrow climate range have a greater indicator value (tolerance-weighted) and also takes the frequency of climate variables in the training data set into account to avoid bias toward high frequency climates (fx-correction) (Liu et al., 2020; Liu et al., 2023). The complete fxTWPALS analysis (training, model selection and application) was performed using the updated fxTWAPLS package (Liu et al., 2020) in R version 4.4.1 (2024-07-05) (R Core Team, 2024). We trained our model based on cover data of the 1387 selected vegetation surveys of the BDM data but excluding both plots with fewer than ten species and species with fewer than two occurrences in the data set, thus relying on a set of 1161 vegetation plots for model calibration. We used binned fx-correction with a bin width of 0.02 using p-spline smoothing, making the fx values nearly independent of the manually provided bin width. The optimal number of components was identified by performing a leave-one-out (LOO) cross-validation and identifying an abrupt increase in the p-value of the root mean square error of prediction (RMSEP; where the p-value assesses the significance of improvement between the current number of components and one less; Table S2). Because cross validation results suggested overfitting when using more than one component for the full fxTWAPLS model, we used the model without tolerance-weighting (fxWAPLS), therefore not downweighting species with wide climatic ranges. In addition to excluding one vegetation plot for the LOO cross-validation procedure, we removed vegetation plots which are geographically (within 50 km horizontal distance, which is the function default of fxTWAPLS) and climatically (within 2% of the full temperature range) close to each other from the training data set to avoid pseudoreplication. Overall compression was assessed by fitting a linear regression to the cross-validation results (fitted values, i.e., fxWAPLS predicted community-inferred temperatures, were used as the response variable and observed plot temperatures as the explanatory variable, n = 1161) and local compression was assessed by locally estimated scatterplot smoothing of the residuals (Figure S1).

Using the trained and validated fxWAPLS regression model, we predicted community-inferred temperatures of all MIREN plots and for each of the three survey years (2012, 2017 and 2022) based on their species composition and species cover values. Additionally, we predicted a second set of community-inferred plot temperature values for the same plots and survey years but based on species composition only, hence removing the effect of cover, by manually setting all cover values to one. This procedure allowed us to assess the impact of species’ abundance changes on community temperature changes. We then fitted linear mixed-effects models (LMMs) to evaluate the rate of change in community-inferred temperature values along the studied elevational gradient and over time between the first survey (2012) and the last resurvey (2022). To explain community-inferred temperatures (i.e. the response variable), plot elevation, survey year and their interaction were included as fixed effects in the model formula with road, transect and plot ID included as nested random intercept terms. As community temperature changes along elevation gradients have been observed to be non-linear, we tested for a potential non-linear response of community-inferred temperature by fitting two models: one with a linear term and one with a second-order polynomial term in the model formula for plot elevation, i.e., poly(elevation, 2). We then compared the two full models (i.e. including the interaction term between survey year and elevation) using a likelihood-ratio test. The model with the better fit included the second-order polynomial term for elevation and was therefore used in the model formula for plot elevation for all further analysis. For comparison purposes, we also evaluated the rate of climate change, in °C/yr, from macroclimatic data by fitting LMMs of mean annual temperature (i.e., the response variable) with plot elevation, year and their interaction term as fixed effects and road and transect as nested random intercept terms. Plot ID was not included as a random intercept term since the macroclimatic data used for this model are identical to the data used for the fxWAPLS model predictions and were only extracted at the transect level and not at the plot level for the MIREN survey data. As for the model using community-inferred temperature as an explanatory variable, we also tested for a non-linear response of mean annual temperature along elevation. Using a second-order polynomial term in the model formula for plot elevation did not improve the fit and we therefore kept the more parsimonious model using a linear term for plot elevation. The significance of model variables was determined by comparing nested models with likelihood-ratio tests. Whenever the interaction term was non-significant, we re-fit the model without the interaction term to extract effect sizes of the additive model. All models were fitted using the lme4 package in R (Bates et al., 2015) and checked for overdispersion and basic model assumptions with diagnostic plots.

To contextualize the number of species gained/lost, we determined the total number of species per plot in each year and fitted a GLMM with species richness as the response variable and a Poisson error distribution. Plot elevation, survey year and their interaction term were included as fixed effects whereas road, transect and plot ID were included as nested random intercept terms. Because species richness might display a hump-shaped relationship along the studied elevational gradient, we tested for a non-linear response of species richness along elevation using the second order polynomial term, as before. Using a second order polynomial term of plot elevation instead of a linear term significantly improved the model fit and was thus retained.

To explore changes in species composition as potential indicators of future community thermophilization, we determined all species that were either gained or lost from a plot between 2012 and 2022, and extracted their species-specific temperatures from the fxWAPLS results.

Then, we fitted a generalized linear mixed-effects model (GLMM), assuming a Poisson error distribution for count data, to explain the number of gained/lost species (i.e., the response variable) with the type of change (gain/loss), plot elevation and their interaction term as fixed effects. To account for pseudoreplication due to the sampling design, we included road, transect and plot ID as nested random intercept terms in the model formula. We found that the final model, determined by comparison of nested models with likelihood-ratio tests, was slightly overdispersed and therefore did a post-fitting adjustment to the coefficient table to apply a quasi-likelihood estimation by multiplying the standard error by the square root of the dispersion factor and recalculating p values (Bolker, 2023).

To further investigate the species gained and lost between 2012 and 2022, we calculated the mean species-specific temperature of gained, lost and consistently-present (i.e. those species present in both 2012 and 2022) species for each plot. We then fitted a LMM to relate the plot mean of gained/lost species (i.e., the response variable) with the plot mean of consistently- present species, including road and transect as nested random intercept terms to account for the nestedness of the data. To assess whether the gained/lost species are a random subset of the consistently-present species, and whether the average species-specific temperatures of the two groups differ, we developed a null model. Our null model randomly samples the real number of gained/lost species from the pool of consistently-present species and calculates their mean species-specific temperatures, re-creating the null expectation that gained/lost species have the same average species-specific temperatures as consistently-present species. We used the thus calculated mean temperatures of the randomly sampled subset of species to fit a linear model between mean species-specific temperatures of randomly sampled species and consistently- present species. We first fit an LMM including road and transect as nested random intercept terms, but decided to use a LM instead because of singularity issues. Repeating the random sampling and modelling process 1000 times allowed us to calculate a trendline and 95% confidence intervals of our null expectation. We then compared the observed trendline for gained/lost species to the null expectation, inferring that gained/lost species deviate significantly in their mean species-specific temperature if the trendline fell outside of the confidence intervals of the null model. To account for the influence of cover on the loss of species (i.e., lower probability of losing dominant species in a plot), the random sampling of potential lost species was weighted by the inverse of species’ cover at the plot level.

Finally, we estimated species’ range shifts by defining the upper and lower range limits of each species (n = 381) as the 90th and 10th percentile of its distribution along the studied elevational gradient in the MIREN data and calculating the difference between 2012 and 2022. We used percentiles instead of the highest/lowest occurrences as a more robust proxy for maximal/minimal elevation in order to avoid influence of extreme outlier occurrences. To compare variation in species-level range shifts along the elevational gradient with the elevation-dependent community changes, we fit two linear models with upper/lower range limit shifts as response variables and species’ range limits during the first survey as explanatory variables.

Range shift observations were weighted by the frequency of occurrence of each species in the complete data set. To account for the limitations of bounded environmental gradients (geometric constraint and regression towards the mean, Iseli et al., 2023; Mazalla & Diekmann, 2022) we created two null models to generate the null expectation of range limit shifts being placed at random along the studied elevational gradient. To do so, our null model randomly assigned the observed range shifts of the upper/lower elevational range limit to new initial elevations along the studied elevational gradient from the pool of surveyed elevations. Potential new initial elevations for each observed range shift were limited to values resulting in final elevations within the surveyed elevational gradient. A weighted linear regression (weighted by the species’ total frequency) of range shifts at the upper/lower elevational range edge (i.e., the response variable) against the randomly sampled initial elevations was then fitted. Random sampling of initial elevations and subsequent model fitting was repeated 1000 times, each time retaining the fitted values of the regression. The fitted values were then used to calculate a 95% confidence interval for the null expectation of randomly occurring shifts along the studied elevational gradient (for more details on the null models, see Iseli et al., 2023). Deviations of the observed real relationships from the null expectation were visually assessed.

## RESULTS

We found a significant increase in community-inferred temperatures that were not weighted by cover values of +0.13°C/year (Χ^2^(1) = 7.545 and p = 0.006 for the effect of year in the LMM). This increase over time did not depend on elevation (Χ^2^ = 1.404 and p = 0.496 for the interaction term between elevation and year in the LMM; Figure 2C, Table S3 and S4).

**Figure 2.**
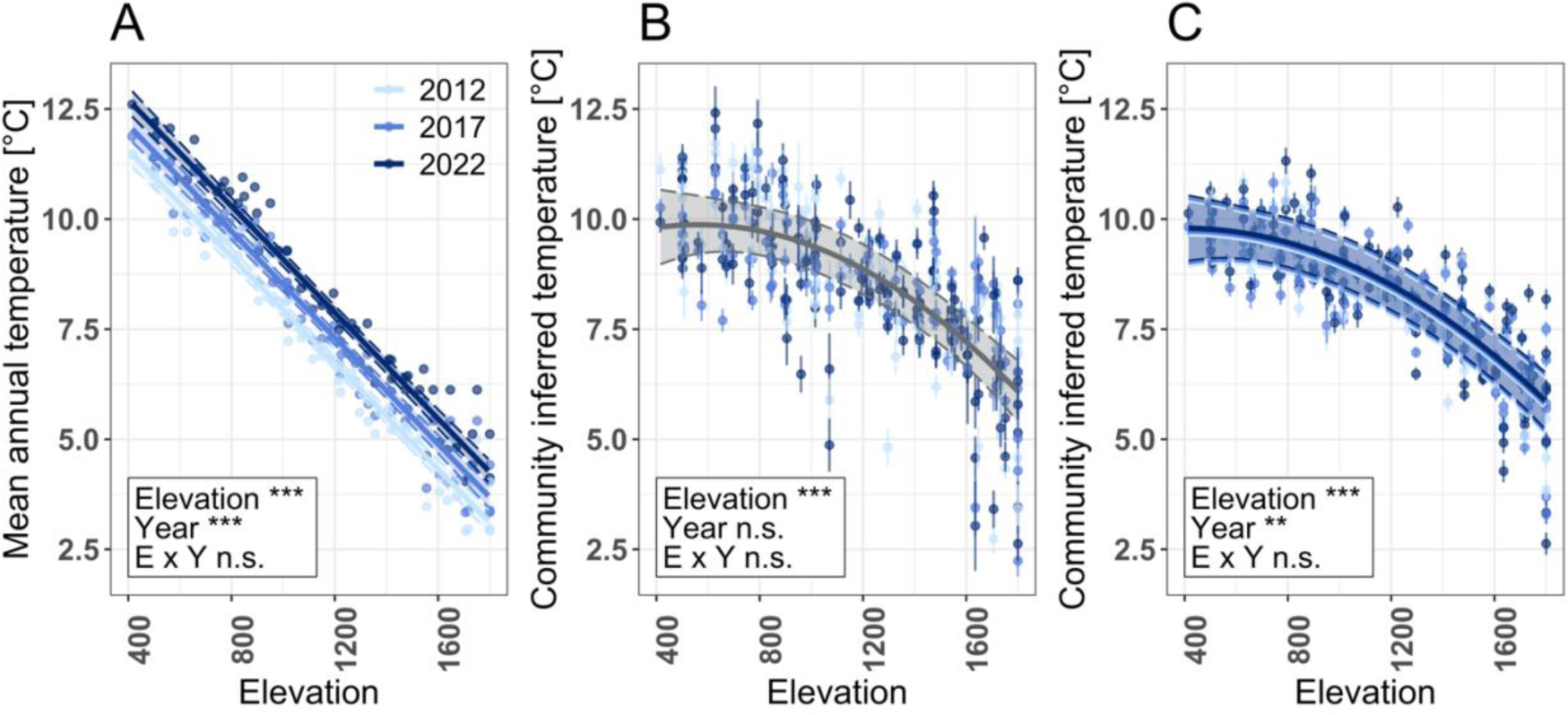
Decreasing community-inferred temperatures along the elevation gradient between 2012 and 2022. (A) Points indicate the rolling average over five years of mean annual temperatures of all 62 repeatedly surveyed MIREN plots in each survey year (2012, 2017 and 2022). Solid trend lines depict the fitted values of the best linear mixed effects model (fixed effects include elevation and survey year, Table S6), dashed lines and shading indicate the 95% confidence interval. (B) and (C) Points and error bars indicate mean community-inferred temperatures and the bootstrapped sample specific error of all 122 repeatedly surveyed MIREN plots in each survey year (2012, 2017 and 2022) as predicted by the fxWAPLS model. (B) Community-inferred temperatures based on species composition and cover values. Solid and dashed grey lines and shading depict the fitted values and 95% confidence interval of the best linear mixed effects model (fixed effects include elevation and the second order polynomial of elevation, Table S4). (C) Community-inferred temperatures based on species composition only (no cover weighting). Solid trend lines depict the fitted values of the best linear mixed effect model (fixed effects include elevation, the second order polynomial of elevation and survey year, Table S4), slim dashed lines and shading indicate the 95% confidence interval. Colors represent survey years (light blue = 2012, medium blue = 2017, dark blue = 2022).

Unweighted community-inferred temperatures significantly decreased with elevation and ranged from 9.79°C at 415 m a.s.l. to 5.76°CC at 1795 m a.s.l. (Χ^2^(2) = 94.868 and p < 0.001 for the effect of elevation and its second order polynomial of elevation in the LMM). In contrast, we found no significant change in community-inferred temperatures between 2012 and 2022 in cover-weighted models (Χ^2^(1) = 0.687 and p = 0.407 for the effect of year in the LMM), and also no elevation-dependence of changes in community-inferred temperatures (Χ^2^ = 0.087 and p = 0.957 for the interaction term between elevation and year in the LMM; Figure 2B, Table S3 and S4). Fitted values of community-inferred temperatures decreased along the studied elevational gradient and ranged from 9.87°C at 565 m a.s.l. to 6.12°C at 1795 m a.s.l. (Χ^2^(2) = 73.676 and p < 0.001 for the effect of elevation and its second order polynomial in the LMM). During the study period, the change in mean annual temperature along the elevational gradient did not vary over time (Χ^2^(1) = 0.378 and p = 0.539 for the interaction term between elevation and year in the LMM; Figure 2A, Table S5 and S6), but mean annual temperature was both highly elevation dependent (Χ^2^(1) = 193.51 and p < 0.001 for the effect of elevation in the LMM) and increased by +1.14°C over the ten year study period (Χ^2^(1) = 453.98 and p = 0.001 for the effect of year in the LMM).

The number of species present in each plot (plot-level species richness) varied both along the elevation gradient and over time, with decreasing species richness at the upper end of the elevational gradient between 2012 and 2022 (significant interaction term between elevation and year in the GLMM; Χ^2^(2) = 7.105, p = 0.029; Figure 3A, Table S7 and S8). The number of species gained and lost per plot depended on the elevation of a plot, with the number of lost species increasing with elevation, while the number of gained species remained constant along the elevational gradient (Χ^2^(1) = 16.718 and p < 0.001 for the interaction term between the effect of the type of change (gained or lost) and elevation in the GLMM; Figure 3B, Table S9 and S10).

**Figure 3.**
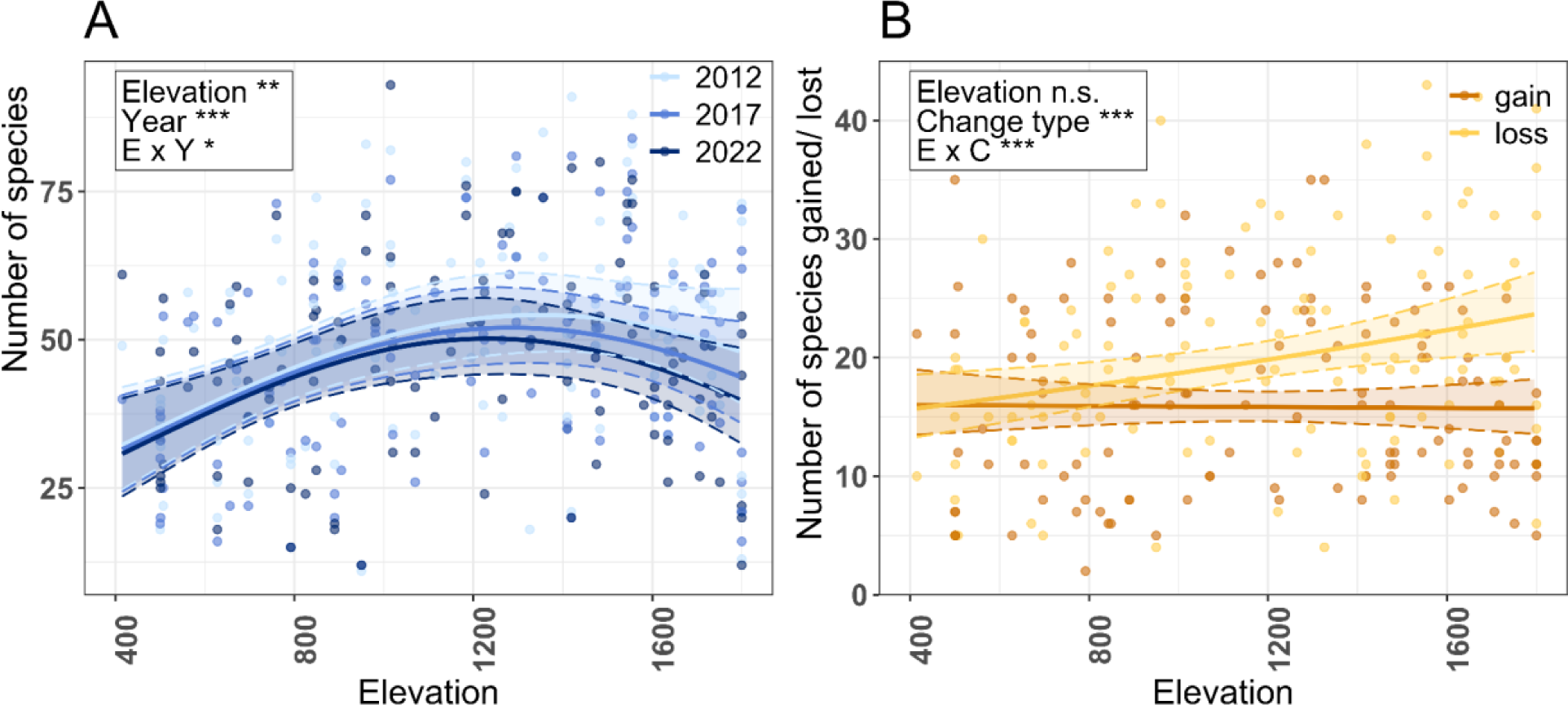
Species richness and number of gained/lost species along the studied elevational gradient. (A) Number of species recorded in each of the 122 repeatedly surveyed MIREN plots for the three survey years. Solid trend lines depict the fitted values of the best generalized linear mixed effects model using a poisson error distribution (fixed effects include elevation, survey year and their interaction, Table S8), slim dashed lines and shading indicate the 95% confidence interval. Colors represent survey years (light blue = 2012, medium blue = 2017, dark blue = 2022). (B) Points indicate the number of gained (orange) and lost (yellow) species in all repeatedly surveyed Swiss MIREN plots. Solid trend lines depict the fitted values of the best generalized linear mixed effects model using a quasi-poisson error distribution (fixed effects include elevation, the type of change — i.e. gain/loss — and their interaction, Table S10), dashed lines and shading indicate the 95% confidence interval of the fixed effects.

To further evaluate the temperature affinity of the gained/lost species, we compared the plot mean of species-specific temperatures of gained/lost species between 2012 and 2022 to the plot mean of species that were consistently present over the same period. The mean temperature of gained species was very similar to the mean of consistently-present species (Figure 4A), with the linear regression falling primarily within the confidence interval of the null model. This indicates that plot means of species-specific temperatures of gained species do not significantly differ from plot means of species-specific temperatures of consistently-present species. In contrast, the observed relationship between the plot means of lost species and the plot means of consistently-present species significantly differed from the null model. The plot means of lost species in warmer plots were significantly lower than plot means of consistently present species (Figure 4B), indicating that especially in warmer plots, species with lower than average species-specific temperatures were lost.

**Figure 4.**
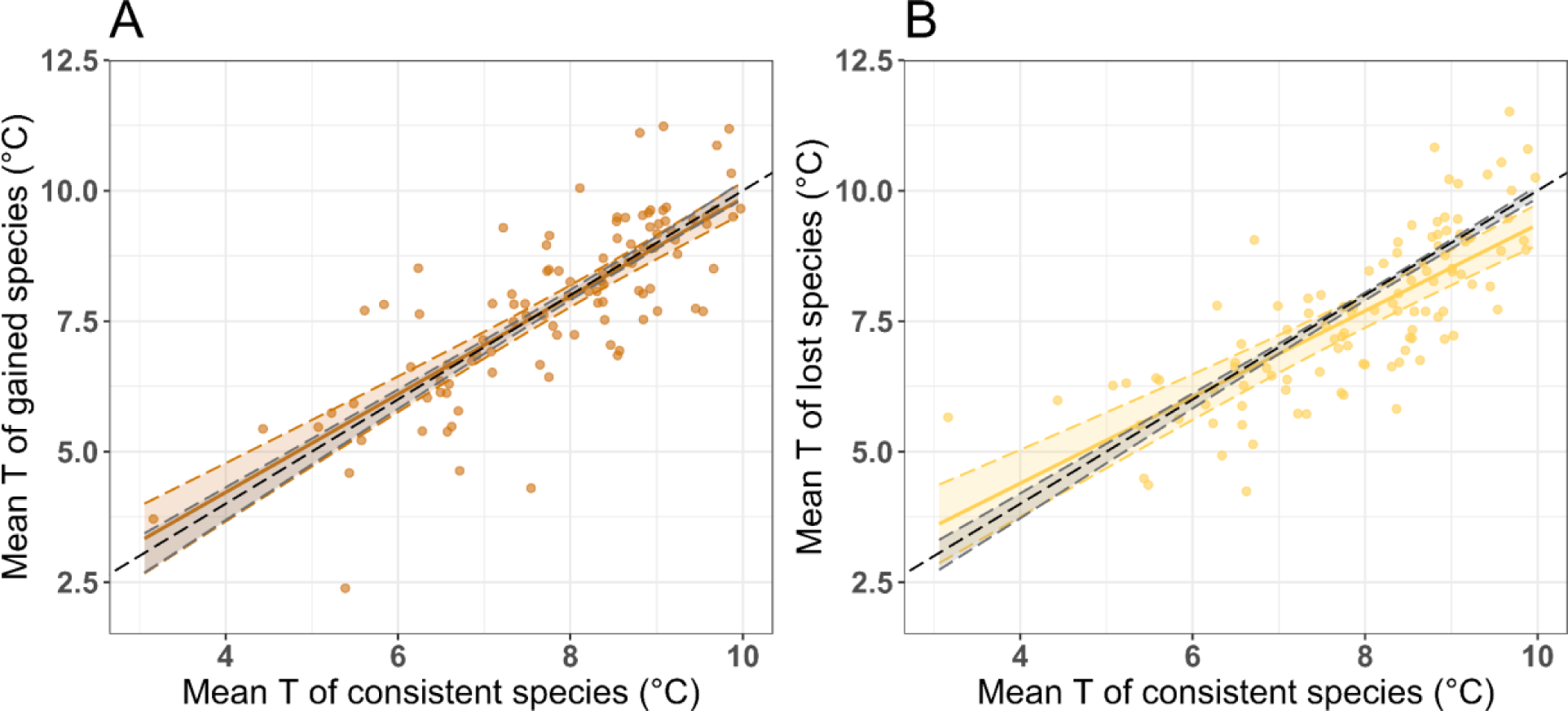
Mean plot temperatures of gained (A) and lost (B) species along the studied elevational gradient between 2012 and 2022. Comparison between plot-level mean of species-specific temperatures extracted from fxTWAPLS model output of species newly occurring (A) or lost (B) in 2022 compared to 2012 versus species that were consistently present over the same time period. The null model (gray dashed lines and shading) is based on random sampling of the number of gained species in the pool of consistently-present species of the same plot, i.e. it shows the expected mean temperatures of gained species if they originated from a species pool identical to the consistently-present species. For the null model of lost species, the random sampling of lost species is inversely weighted by the species’ cover value on the plot level. Points indicate the 62 repeatedly surveyed MIREN permanent plots. Coloured solid lines depict model fits of a linear mixed effects model (orange = gained species, yellow = lost species), slim dashed lines and shading 95% confidence intervals and black dashed line shows the 1:1 line.

At the species level, species shifted their upper range limits upwards more than expected by chance in the lower two thirds of the elevational gradient, while species with a high initial range limit showed no or non-significant range limit shifts (Figure 5A). After accounting for the effect of elevation on upper range limit shifts by evaluating mean shifts in the middle of the elevational gradient, we observed significant upward shifts of the upper range limits by on average 88.9 ± 16.12m (fitted value ± standard error) between 2012 and 2022 (Figure 5A). In contrast, species’ lower range limits shifted downwards on average by 34.64 ± 11.69m (fitted value ± standard error) over the same period, but not more than expected by chance at either end of the studied elevational gradient (Figure 5B).

**Figure 5.**
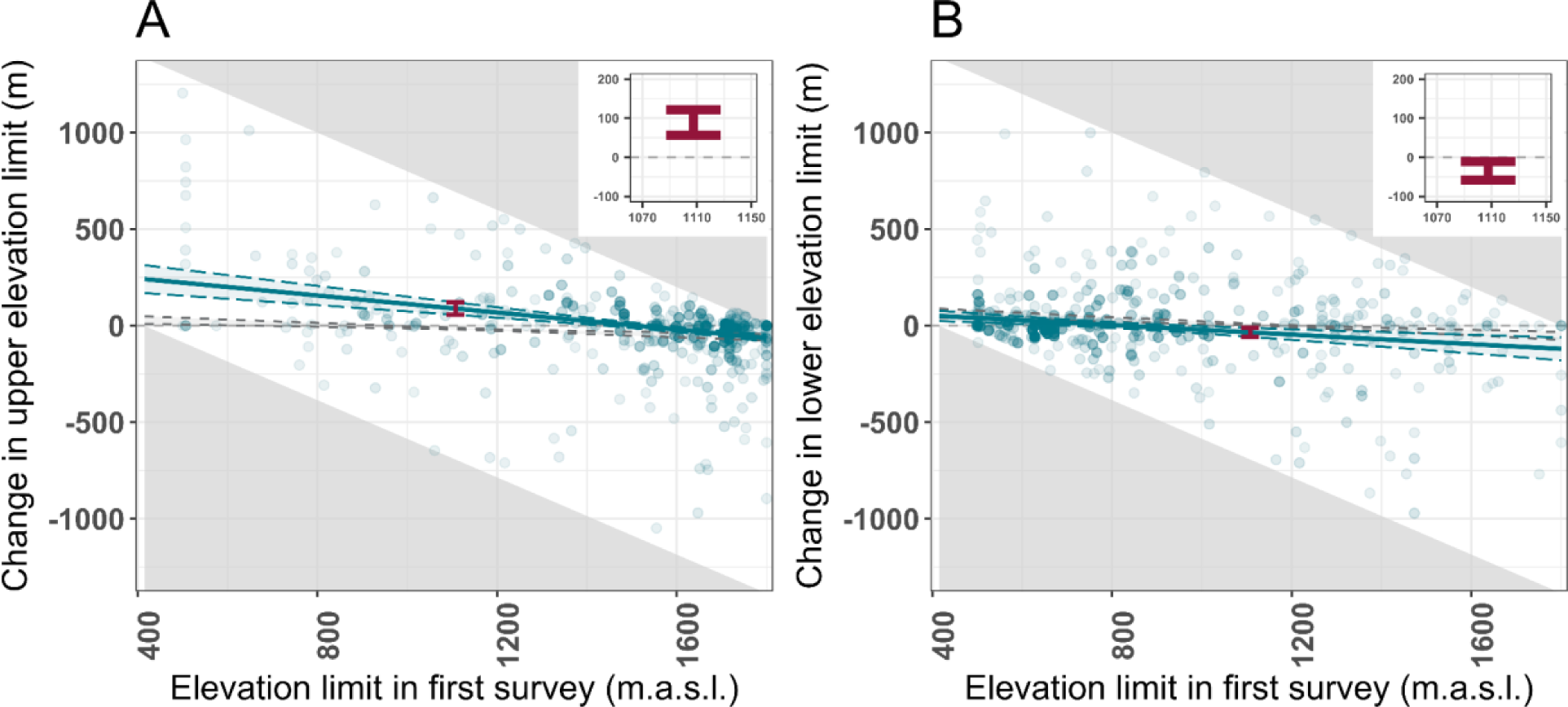
Change in upper (A) and lower (B) elevational limits between 2012 and 2022 in relation to a null model. Points depict all species recorded in the Swiss MIREN surveys at least once both in 2012 and 2022. Solid and slim dashed teal lines and shading indicate the fitted relationships and 95% confidence intervals of the relationships. Red error bars highlight the range limit shift and 95% confidence interval in the middle of the studied elevational gradient, which is also shown in the inset. Gray triangles mark zones where range shifts are unobservable due to constraints imposed by the limits of the surveyed elevational gradient. Gray dashed lines indicate the 95% confidence interval of the null expectation between original upper/lower range limit and change between 2012 and 2022.

## DISCUSSION

Our analyses of community thermophilization in the Swiss Alps — using vegetation survey data along three mountain roads between 2012 and 2022 — reveal a complex relationship between changes in species richness, species’ range shifts and abundance changes. Contrary to our hypothesis, combining species distribution and abundance changes at the community level by calculating community-inferred temperatures did not yield a clear signal of community warming over the relatively short study period of 10 years. While we did detect a significant thermophilization signal of +0.13 °C over this time period when only considering changes in species presence-absence, we found no significant increase in community-inferred temperatures when incorporating species’ cover values. Additionally, we observed pronounced species-level range shifts at the lower end of the studied elevation gradient. These findings can be explained by species shifting their ranges within elevation bands that have similar temperatures to their own species-specific temperature (i.e., the species-specific temperatures of gained/lost species did not differ from plot averages) and by the low abundance of gained and lost species differing from plot averages. Overall, our results indicate that while observable responses to climate warming are happening at both the community and species level, community changes are concealed when species’ covers are taken into account.

Our result of significant increases in community-inferred temperatures over time based on presence-absence data align with previous findings on Himalayan mountain summits (Hamid et al., 2020; Verma et al., 2023), in understory plants and highland temperate forests in France (Bertrand et al., 2011; Martin et al., 2019; Richard et al., 2021), in tropical forests in South America (Freeman et al., 2021), alpine plant communities in Europe (Gottfried et al., 2012), and across Switzerland as a whole (Kiebacher et al., 2023). Additional non-significant increases have been observed on Andean summits and in Andean tropical forests (Cuesta et al., 2023; Duque et al., 2015; Fadrique et al., 2018), European and Canadian forests (Savage & Vellend, 2015; Zellweger et al., 2020), tropical trees in Costa Rica (Feeley et al., 2013) and alpine plants in Austria (Lamprecht et al., 2018), whereby there are no immediately apparent differences such as study duration that might explain differences between significant and non-significant trends . Reported increases in community-inferred temperatures range from 0.001°C/year (Zellweger et al., 2020) to 0.027°C/year (Duque et al., 2015) and there is no obvious trend for greater increases in studies reporting significant thermophilization rates or in studies focusing on longer time periods. Although the majority of other thermophilization studies found increasing community-inferred temperatures, slight decreases have also been observed in smaller scale analyses (e.g. Verma et al. (2023) at a Northern aspect of a Himalayan mountain summit, Kiebacher et al. (2023) in lowland managed grasslands). Most frequently the observed community warming has been explained by upward range shifts and increasing species richness of warm-adapted species (Bertrand et al., 2011; Cuesta et al., 2023; Hamid et al., 2020; Kiebacher et al., 2023; Lamprecht et al., 2018; Richard et al., 2021; Verma et al., 2023), loss of cold-adapted species (Duque et al., 2015; Feeley et al., 2013; Lamprecht et al., 2018) and changes in species abundances (Bertrand et al., 2011; Cuesta et al., 2023; Fadrique et al., 2018; Feeley et al., 2013; Hamid et al., 2020; Lamprecht et al., 2018; Martin et al., 2019; Verma et al., 2023). In contrast to other studies including an elevational gradient (Bertrand et al., 2011; Fadrique et al., 2018; Hamid et al., 2020; Kiebacher et al., 2023; Savage & Vellend, 2015; Verma et al., 2023), we did not find any elevation dependency of community thermophilization. However, the elevational gradient we studied, spanning 415–1800 m a.s.l., was relatively short compared to elevational gradients covered in former studies (Bertrand et al., 2011; Fadrique et al., 2018; Kiebacher et al., 2023), and its upper elevational limit was low compared to Hamid et al. (2020) and Verma et al. (2023). Our comparatively short and low elevational gradient therefore did not include (high) alpine communities, which could be expected to react disproportionately strongly to climate warming due to elevation dependent warming (Pepin et al., 2015).

Plant community changes are in general lagging behind the pace of change expected based on the current rate of climate warming (Alexander et al., 2018). This lag in the response of plant communities to climate warming is often expressed as a slower increase in community temperatures compared to macroclimatic or even microclimatic temperatures, as is the case in our study (i.e., the increase of mean annual temperature being +1.14°C/ decade vs. the increase in unweighted community-inferred temperature of +0.13°C/decade). Such thermal lags of plant communities have been detected across a broad range of ecosystems over the last decade (Bertrand et al., 2011; Bertrand et al., 2016; Freeman et al., 2021; Lenoir et al., 2020; Richard et al., 2021) and lags have been found to accumulate over time (Pacheco-Riaño et al., 2023). Variation in thermal lags between ecosystems has been related to elevational differences (Bertrand et al., 2011; Freeman et al., 2021), the historical temperature of study areas (Bertrand et al., 2016), temperature-change velocity (Pacheco-Riaño et al., 2023) and forest structure (Richard et al., 2021). However, although such broad scale analyses of community temperatures and thermal lags give insight into a specific component of community change, i.e., whether the community-weighted mean of all species-specific temperatures within a community changes, they potentially overlook changes in community composition and species distributions which are not reflected in this metric. Our study highlights this limitation by reporting highly significant species-level range shifts in the lower two thirds of the studied elevational gradient and significantly decreasing species richness at the upper end of the studied elevational gradient, while at the same time only the community-inferred temperatures calculated without incorporating species’ cover values show a significant increase. Theoretically, lower-elevation species shifting their upper range limit to higher elevation can be assumed to be at their cold- edge and therefore have a higher species-specific temperature than the communities they colonize; they could therefore be expected to increase community-inferred temperatures.

Similarly, decreasing species richness at high elevations could be expected to lead to community warming, too, assuming cold-adapted species from high elevations are more likely to react negatively to warming temperatures and be lost from survey plots compared to species with a warmer species-specific temperature. Based on our analysis on the effect of abundance changes on community thermophilization rates and the temperature affinity of gained/lost species, we present two reasons as to why the observed species-level range shifts and changes in species richness didn’t translate into increasing cover-weighted community-inferred temperatures.

Firstly, the difference between the significant +0.13°C/decade increase in community-inferred temperatures calculated without cover-weighting and the non-significant change for cover-weighted community-inferred temperatures indicates that including species’ cover masked the impact of gained/lost species rather than reinforcing community warming in our study. Even after successfully establishing within a new community, species are likely to be found at a low abundance when they first arrive. Population growth after establishment is limited by the life cycle of a species and especially long-lived perennials such as trees and other woody species often take multiple years or even decades to reach maturity and increase in cover (Svenning & Sandel, 2013). Lag times between arrival and population expansion can be further extended by abiotic stresses or competition (Svenning & Sandel, 2013). Consequently, given that cover-weighted community properties are heavily influenced by more dominant species, there will be a lag between species joining a community and them causing observable changes in community properties. Likewise, species that are lost from one survey time point to the next will likely already have been at low abundance at the previous survey time point, contributing little to cover-weighted community-inferred temperatures. Indeed, in our study, species that were lost between 2012 and 2022 showed significantly lower cover values (0.27 ± 1.16, mean ± sd) than consistently-present species (1.03 ± 3.46, mean ± sd, Student’s t-test, t(81468) = 65.96, p < 0.001). Low cover values are most likely one of the reasons why we don’t see cover-weighted community thermophilization at the lower (warmer) end of the elevational gradient, even though here, lost species on average exhibited lower species-specific temperatures than consistently- present species. Detecting species loss during the relatively short study period of ten years supports recent findings that local extinction lags can often be short (Nomoto & Alexander, 2021) and can therefore be an important driver of community change. However, although our analysis revealed a tendency for downward shifts of lower range limits at the upper end of the studied elevational gradient (this observation is based on a relatively low number of species at the edge of the studied elevational gradient and should be treated with caution), we found no significant upwards shift of lower range limits at the lower end of the gradient. The loss of species with cooler species-specific temperatures compared to the consistently-present species at the lower end of the studied elevational gradient can therefore not be unambiguously interpreted as local extinction events due to high-elevation species retracting their lower range limit to higher elevations.

Secondly, we identified the species-specific temperatures of gained/lost species as another reason for the non-significant increase in cover-weighted community-inferred temperatures. Examining the species-specific temperatures of species that are gained/lost over the study period shows that mean species-specific temperatures of gained species along the whole elevational gradient, and lost species in the upper part of the elevational gradient, did not differ from the plot-level averages of consistently-present species. Consequently, if species establishing or being lost from a plot don’t deviate from the rest of the community in their temperature affinity, changes in the community composition have minimal influence on community-inferred temperatures. The similarity of species-specific temperatures between gained and consistently-present species could indicate that although species are dispersing across the elevational gradient, they are only covering relatively short distances and are merely moving within their current thermal band. The effect of such dispersal lags (Alexander et al., 2018) could be amplified in regions and ecosystems with poorly connected pools of species with warmer temperature affinities, as suggested by Bertrand et al. (2011), Bertrand et al. (2016) and Savage and Vellend (2015) for forest communities. To reach communities at the lower end of an elevational gradient, warm-adapted species are required to disperse across longer distances along latitudinal gradients and might be impeded by barriers such as mountain chains (Lawlor et al., 2024), as is the case in Switzerland due to the Alps (Kiebacher et al., 2023). As the lower limit of our studied elevational gradient is the bottom of a mountain valley within the Swiss Alps, dispersal lags could be an important reason for why significant range shifts in the lower two thirds of the elevational gradient did not result in increasing cover-weighted community-inferred temperatures. Alternatively, the influence of dispersing warm-adapted species could be limited due to establishment lags, meaning that although species reach new communities in cooler conditions, they are not yet able to establish.

Overall, our results suggest that especially in systems mainly consisting of locally common species with broad and similar temperature affinities, cover-weighted community-inferred temperatures will fail to detect small, but still potentially important, changes in community composition. However, community-inferred temperatures are only one way to characterize shifts in community composition following climate change. For example, Savage and Vellend (2015) include alternative community properties such as the community precipitation index and the community light index together with the community temperature index in their analysis of community-level consequences of range shifts on community warming. This approach might be broadened to include species functional traits related to resource capture and use, which might generate a more differentiated picture of the effect of climate change on plant communities and facilitate a more mechanistic understanding of community composition changes. Additionally, cover-weighted community metrics bear the risk of being highly affected by the environmental conditions of the survey year. Dry summers and water deficits in particular can cause temporary cover decreases in susceptible plants, even if all individuals survive the dry period and recover. Using species abundance (i.e., the number of individuals per species) instead of cover might reduce the impact of short-term unfavourable environmental conditions, but is much more time-intensive to record and only represented in broad classes in many available longer-term vegetation survey datasets (both datasets used in this study record abundance in three abundance classes, which also differ between the datasets). This sensitivity to the environmental conditions in the survey year together with potential observer effects decreases with an increasing number of survey time points when survey year effects are overridden by long-term trends (Lawlor et al., 2024). Indeed, if – unlike in the analyses presented earlier – the change in community-inferred temperatures is modelled with year as a factor rather than a continuous variable, we found larger changes in community-inferred temperatures between 2012 and 2017 than between 2012 and 2022, implying that community warming should not be treated as a linear process (Table S11).

In conclusion, our findings show that concentrating on cover (or abundance) weighted community thermophilization alone as an indicator of community thermophilization does not sensitively capture short-term (i.e. over a 10 year period) community responses to climate change. Focusing on common species by cover-weighting and only considering a single community characteristic such as community-inferred temperature without the context of species-level changes can result in a misleading interpretation of relatively static communities, resulting in monitoring and management recommendations not optimally matched to the real circumstances. However, the detection of species-level range shifts requires a relatively comprehensive dataset and even surveys explicitly designed to capture range shifts — such as that conducted by the MIREN network — are fairly labor intensive and might not be feasible for regional small-scale projects. Consequently, we recommend the use of community thermophilization based on presence-absence data only for an efficient early detection of climate change impacts on plant communities.

## Acknowledgements

We thank Fiona Schwaller and the rest of the Swiss MIREN team for their contribution to the data collection. The Swiss Federal Office for the Environment (FOEN) kindly provided the BDM data. We thank the dedicated team who conducted the fieldwork for the Swiss Biodiversity monitoring program. The survey data can be requested from Biodiversity Monitoring Switzerland (www.biodiversitymonitoring.ch). This research was funded by the 2019–2020 BiodivERsA joint call for research proposals, under the BiodivClim ERA-Net COFUND programme, with the funding organisations the Swiss National Science Foundation (Grant No. 20BD21_193809), Innovation Fund Denmark, the Department of Science and Innovation Republic of South Africa, the Research Council of Norway, the Swedish Research Council for Environment, Agricultural Sciences and Spatial Planning and the German Research Foundation. J.L. was supported by the synthesis center (CESAB) of the Fondation pour la Recherche sur la Biodiversité (FRB; www.fondationbiodiversite.fr) via the project FRAGSHIFTS funded by the French Ministère de la Transition Écologique (MTE), the Office Français de la Biodiversité (OFB) and the FRB.

## Author contributions

J.A., E.I. and J.L. designed the study, C.B. and N.D.Z. collected and managed data. E.I. analysed the data and drafted the manuscript with support and feedback from J.A. and J.L.. All authors edited and commented on the paper.

## Data availability statement

The complete MIREN data set up to 2017 is published and available on Zenodo (https://doi.org/10.5281/zenodo.7495407), the Swiss data of the MIREN survey from 2022 is provided for review purposes and will be published latest on publication of this manuscript. The Swiss Biodiversity Monitoring data were provided by Biodiversity Monitoring Switzerland (BDM, www.biodiversitymonitoring.ch) on inquiry, but restrictions apply to the data availability and they are not publicly available. The climate data used is extracted from and available on CHELSA cruts (https://chelsa-climate.org/chelsacruts/) and TerraClim (www.climatologylab.org/terraclimate.html).

