## SupplementaryMaterial for "Early signs of plant community responses to climate warming along mountain roads in Switzerland"

**Supplementary material**

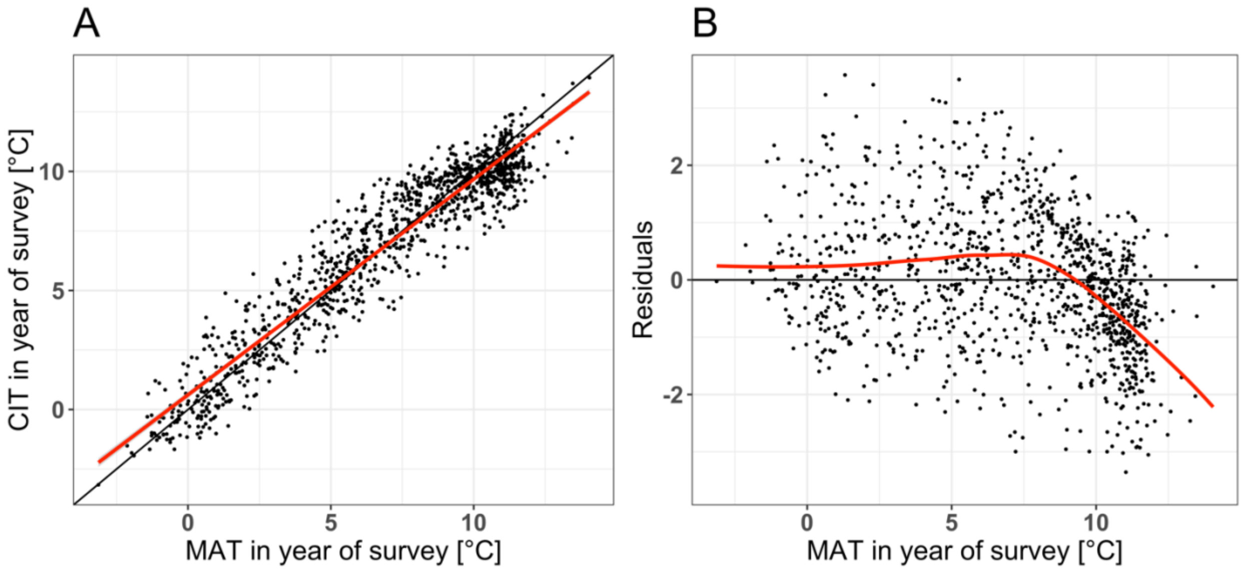


**Figure S1. Control plots to assess artificial compression of fxWAPLS model results.** Community-inferred temperatures predicted by fxWAPLS using the last significant number of components (A) and their residuals (B) related to the observed mean annual temperatures of the Swiss Biodiversity Monitoring training data set. The black line indicates the 1:1 line (A) and the zero line (B), the red line the linear regression between the described variables.

**Table S1.** Conversion of cover classes between the Swiss Biodiversity Monitoring data (BDM), the MIREN data collected based on the 2016 MIREN manual (MIREN16) and the MIREN data collected based on the 2021 MIREN manual (MIREN21).

| BDM cover classes | BDM cover values | BDM used values | MIREN cover classes | MIREN21 cover values | MIREN21 used values | MIREN16 cover values | MIREN16 used values |
|---|---|---|---|---|---|---|---|
| r | < 0.1 | 0.1 | 1 | < 0.1 | 0.1 | <1 | 0.1 |
| + | 0.1 - 1 | 0.5 | 2 | 0.1 - 1 | 0.5 | 1 - 5 | 3 |
| 1 | 1 - 5 | 3 | 3 | 1 - 5 | 3 | 6 - 25 | 15 |
| 2a | 5 - 15 | 10 | 4 | 5 - 10 | 10 | 26 - 50 | 37.5 |
| 2b | 15 - 25 | 20 | 5 | 10 - 25 | 20 | 51 - 75 | 62.5 |
| 3 | 25 - 50 | 37.5 | 6 | 25 - 50 | 37.5 | 76 - 95 | 85 |
| 4 | 50 - 75 | 67.5 | 7 | 50 - 75 | 67.5 | 96 - 99 | 97.5 |
| 5 | 75 - 100 | 87.5 | 8 | 75 - 100 | 87.5 | 100 | 100 |

**Table S2.** Results from the pseudo-removed leave-one out cross validation used to determine the last significant number of components for the fxTWAPLS (with tolerance weighting) and fxWAPLS (without tolerance weighting) models used to predict community-inferred temperatures. Geographically and climatically close plots were excluded to avoid pseud-replication. ncomp is the number of components, RMSEP is the root mean square error of prediction, ΔRMSEP the percent change in RMSEP when using one component less than the current number. p assesses whether using one component more than the current number is significantly better, which is used to select the last significant number of components without overfitting (indicated in bold). $b_0$ and $b_1$ are the intercept and slope of the linear regression of the fitted values (predicted community-inferred temperatures) related to the training data (mean annual temperatures), $b_0$.se and $b_1$.se their standard error. The closer to one the slope, the less the overall compression (Liu at al. 2020).

| | ncomp | $R^2$ | RMSEP | ΔRMSEP | p | $b_0$ | $b_1$ | $b_0$.se | $b_1$.se |
|---|---|---|---|---|---|---|---|---|---|
| fxWAPLS | 1 | 0.8594 | 1.4112 | -62.4644 | 0.0010 | 0.8894 | 0.8627 | 0.0812 | 0.0102 |
| | 2 | 0.8672 | 1.3717 | -2.8018 | **0.0040** | 0.8146 | 0.8752 | 0.0797 | 0.0101 |
| | 3 | 0.8704 | 1.3667 | -0.3616 | 0.3906 | 0.5547 | 0.9166 | 0.0823 | 0.0104 |
| | 4 | 0.8601 | 1.4225 | 4.0846 | 1.0000 | 0.6195 | 0.9127 | 0.0857 | 0.0108 |
| | 5 | 0.8548 | 1.4496 | 1.9012 | 0.9850 | 0.6513 | 0.9094 | 0.0872 | 0.0110 |
| fxTWAPLS | 1 | 0.8807 | 1.3050 | -65.2887 | **0.0010** | 0.5688 | 0.9076 | 0.0777 | 0.0098 |
| | 2 | 0.8818 | 1.2952 | -0.7551 | 0.2517 | 0.6639 | 0.8951 | 0.0762 | 0.0096 |
| | 3 | 0.8787 | 1.3164 | 1.6413 | 0.9341 | 0.5770 | 0.9058 | 0.0783 | 0.0099 |
| | 4 | 0.8732 | 1.3496 | 2.5201 | 0.9970 | 0.5492 | 0.9136 | 0.0810 | 0.0102 |
| | 5 | 0.8641 | 1.3989 | 3.6498 | 1.0000 | 0.5738 | 0.9071 | 0.0837 | 0.0106 |

**Table S3.** Linear mixed-effect model results of the relationship between community-inferred temperature and plot elevation (second-order polynomial relationship), year (continuous) and their interaction. Road, transect and plot ID are included as random effects to account for the nested structure of the data. Models were fitted with community-inferred temperatures based on fx corrected weighted averaging partial least square models with and without incorporating cover values. Sample size N = 366 plots along three roads at three timepoints (2012/ 2017/ 2022).

| | Community-inferred temperature (cover-weighted) | | | Community-inferred temperature (not cover-weighted) | | |
|---|---|---|---|---|---|---|
| Fixed Effects | $X^2$ | df | P | $X^2$ | df | P |
| poly(Elevation, 2) (E) | 73.676 | 2 | **< 0.001** | 94.868 | 2 | **< 0.001** |
| Survey year (Y) | 0.687 | 1 | 0.407 | 7.545 | 1 | **0.006** |
| E x Y | 0.087 | 2 | 0.957 | 1.404 | 2 | 0.496 |

**Table S4.** Linear mixed-effect model results of the best models explaining the relationship between community-inferred temperature, plot elevation (second-order polynomial relationship), year (continuous). Road, transect and plot ID are included as random effects to account for the nested structure of the data. Models were fitted with community-inferred temperatures based on fx corrected weighted averaging partial least square models with and without incorporating cover values. Sample size N = 366 plots along three roads at three timepoints (2012/ 2017/ 2022).

| | Community-inferred temperature (cover-weighted) | | | Community-inferred temperature (not cover-weighted) | | |
|---|---|---|---|---|---|---|
| | Estimates | CI | p | Estimates | CI | p |
| (Intercept) | 8.49 | 7.99 – 9.00 | **<0.001** | 24.81 | -13.96 – 63.59 | 0.209 |
| Survey year | | | | -0.01 | -0.03 – 0.01 | 0.408 |
| Plot elevation | -23.05 | -26.92 – -19.18 | **<0.001** | -23.05 | -26.92 – -19.18 | **<0.001** |
| poly(Plot elevation, 2) | -6.73 | -10.58 – -2.89 | **0.001** | -6.73 | -10.58 – -2.89 | **0.001** |
| Marginal $R^2$ / Conditional $R^2$ | 0.511 / 0.812 | | | 0.511 / 0.811 | | |

**Table S5.** Linear mixed-effect model results of the relationship between mean annual temperature and plot elevation, year (continuous) and their interaction. Road and transect are included as random effects to account for the data's nested structure. Sample size N = 186 transects along three roads at three timepoints (2012/ 2017/ 2022).

| Fixed Effects | $X^2$ | df | P |
|---|---|---|---|
| Plot elevation (E) | 193.51 | 1 | **< 0.001** |
| Survey year (Y) | 453.98 | 1 | **< 0.001** |
| E x Y | 0.378 | 1 | 0.539 |

**Table S6.** Linear mixed-effect model results of the best model explaining the relationship between mean annual temperature, plot elevation and year (continuous). Road and transect are included as random effects to account for the data's nested structure. Sample size N = 186 transects along three roads at three timepoints (2012/ 2017/ 2022).

| | *Estimates* | *CI* | *p* |
|---|---|---|---|
| (Intercept) | -216.07 | -222.74 – -209.39 | **<0.001** |
| Elevation | -0.01 | -0.01 – -0.01 | **<0.001** |
| Year | 0.11 | 0.11 – 0.12 | **<0.001** |
| Marginal $R^2$ / Conditional $R^2$ | 0.955 / 0.999 | | |

**Table S7.** Generalized mixed-effect model results of the relationship between species number per plot (species richness) and the variables year, plot elevation (second-order polynomial relationship) and their interaction, assuming a Poisson error distribution. Road, transect and plot ID are included as random effects to account for the data's nested structure. Sample size N = 366 plots along three roads at three timepoints (2012/ 2017/ 2022).

| Fixed Effects | $X^2$ | df | P |
|---|---|---|---|
| poly(elevation, 2) (E) | 9.261 | 2 | **0.01** |
| Year (Y) | 22.081 | 1 | **< 0.001** |
| E x Y | 7.105 | 2 | **0.029** |

**Table S8.** Generalized mixed-effect model results of the best model explaining the relationship between species number per plot and the variables year, plot elevation (second-order polynomial relationship) and their interaction, assuming a Poisson error distribution. Road, transect and plot ID are included as random effects to account for the data's nested structure. Sample size N = 366 plots along three roads at three timepoints (2012/ 2017/ 2022).

| | *Estimates* | *CI* | *p* |
|---|---|---|---|
| (Intercept) | -513.93 | -1600.83 – 572.98 | 0.353 |
| Year | 0.27 | -0.26 – 0.81 | 0.318 |
| Elevation | 1.19 | 0.32 – 2.05 | **0.007** |
| Year × Elevation | -0.00058 | -0.00101 – -0.00016 | **0.008** |
| Marginal $R^2$ / Conditional $R^2$ | 0.053 / 0.832 | | |

**Table S9.** Generalized mixed-effect model results of the relationship between changes in species number per plot and the variables type of change (gain vs. loss), plot elevation and their interaction, assuming a Poisson error distribution. Road, transect and plot ID are included as random effects to account for the data's nested structure. Sample size N = 244 counts of gained/ lost species along three roads at three timepoints (2012/ 2017/ 2022).

| Fixed Effects | $X^2$ | df | P |
|---|---|---|---|
| Plot elevation (E) | 3.037 | 1 | 0.081 |
| Type of change (C) | 58.018 | 1 | **< 0.001** |
| E x C | 16.718 | 1 | **< 0.001** |

**Table S10.** Generalized mixed-effect model results of the best model explaining the relationship between changes in species number per plot and the variables type of change (gain vs. loss), plot elevation and their interaction, assuming a Poisson error distribution. Road, transect and plot ID are included as random effects to account for the data's nested structure. Sample size N = 244 counts of gained/ lost species along three roads at three timepoints (2012/ 2017/ 2022).

| | *Log-Mean* | *CI* | *p* |
|---|---|---|---|
| (Intercept) | 2.78 | 2.54 – 3.02 | **<0.001** |
| turnover type [loss] | -0.15 | -0.33 – 0.03 | 0.109 |
| Elevation | -0.00001 | -0.00021 – 0.00018 | 0.888 |
| turnover type [loss] × Elevation | 0.00031 | 0.00017 – 0.00045 | **<0.001** |
| Marginal $R^2$ | 0.264 | | |

**Table S11.** Linear mixed-effect model results of the relationship between community-inferred temperature and plot elevation (second-order polynomial relationship), year (factor) and their interaction. Road, transect and plot ID are included as random effects to account for the nested structure of the data. Models were fitted with community-inferred temperatures based on fx corrected weighted averaging partial least square models with and without incorporating cover values. Sample size N = 366 plots along three roads at three timepoints (2012/ 2017/ 2022).

| | Community-inferred temperature (cover-weighted) | | | Community-inferred temperature (not cover-weighted) | | |
|---|---|---|---|---|---|---|
| Fixed Effects | $X^2$ | df | P | $X^2$ | df | P |
| poly(Elevation, 2) (E) | 73.676 | 2 | **< 0.001** | 94.868 | 2 | **< 0.001** |
| Survey year (Y) | 2.249 | 2 | 0.325 | 8.4116 | 2 | **0.015** |
| E x Y | 19.198 | 4 | **0.001** | 15.084 | 4 | **0.005** |
